# Gradient of Wnt signaling facilitates Mef2 heterogeneity and limits commitment of the developmental muscle progenitor pool

**DOI:** 10.1101/2025.07.17.665417

**Authors:** Laura Schütze, Maria Jelena Musillo, Bilyana Popova, Natalja Engel, Verena Holzwarth, Milena Lucia Hild, Anna Geissdorf, Jennifer Nadine Gross, Ingrid Lohmann, Josephine Bageritz

**Affiliations:** Heidelberg University, COS, Stem Cell Niche Heterogeneity, 69120 Heidelberg Germany; Heidelberg University, COS, Developmental Biology, 69120 Heidelberg Germany

**Author notes:** corresponding author correspondence: Dr. Josephine Bageritz. contributed equally (Laura Schütze, Maria Jelena Musillo).

## Abstract

During skeletal muscle development, the timing and extent of lineage commitment towards differentiation must be coordinated to ensure proper tissue formation while preserving undifferentiated progenitors for adult stem cell function. This balance requires spatial and temporal regulation of cell fate and transcriptional regulators to act as key mediators of lineage progression. The transcription factor Mef2 is a key activator of myogenic differentiation across vertebrates and invertebrates. However, the mechanisms that spatially restrict *Mef2* expression within the developing niche to limit muscle progenitor (MP) commitment towards differentiation remains less well characterized. Using the developmental flight muscle progenitor niche in *Drosophila*, a system that parallels vertebrate myogenesis, we investigated the transcriptional and spatial regulation of MP fate at a developmental timepoint when *Mef2* expression begins to rise but precedes overt differentiation.

We identified a spatially distinct subpopulation of MPs with low *Mef2* expression and elevated Wnt/β-catenin signaling. Within the broader MP pool, graded Wnt activity emerged as a key source of Mef2 heterogeneity: high Wnt activity led to strong repression of Mef2 via Armadillo/β-catenin and TCF-dependent regulation. Moderate Wnt activity, meanwhile, not only repressed *Mef2* expression but also sustained expression of *zfh1*, a conserved transcription factor linked to MP maintenance and adult muscle stem cell identity. Contrary to its well-established role in promoting myogenesis, Wnt/β-catenin signaling in this early spatial context instead promotes a less committed state, leading to a subset of MPs with particular low Mef2 level. These findings highlight greater spatial complexity within the developing muscle progenitor niche than previously recognized, with potential implications for conserved strategies of muscle stem cell regulation across species.

## Introduction

Muscle progenitors (MPs) play a critical role in establishing the adult muscle stem cell pool, known as satellite cells, which are essential for tissue homeostasis and lifelong regenerative capacity. While a subset of these developmental progenitors contributes to muscle formation through fusion, others are selected to remain undifferentiated and persist as stem cells. In both vertebrates and invertebrates, MPs follow a defined myogenic lineage progression: mononucleated progenitor cells differentiate into myoblasts/myocytes, which then fuse to form myotubes and mature into myofibers (reviewed in Bothe & Baylies, 2016; reviewed in Schmidt et al., 2019). The balance between differentiation and maintenance of an undifferentiated state is tightly regulated by key transcription factors. PAX7 in vertebrates and its *Drosophila* functional analog Zfh1 actively maintain muscle progenitors in an undifferentiated state by promoting stemness and preventing premature differentiation (Boukhatmi & Bray, 2018; Relaix et al., 2005). Zfh1 exists in two isoforms, a long and a short form, both contribute to this maintenance function (Boukhatmi & Bray, 2018). Together, these activities ensures that a pool of progenitors is preserved until they receive the appropriate environmental cues to activate the differentiation program. The undifferentiated state of muscle progenitors is further stabilized by transcriptional repression and post-transcriptional inhibition of key myogenic effectors that promote differentiation, including members of the MyoD family of basic helix–loop–helix (bHLH) transcription factors, also known as myogenic regulatory factors (MRFs), and the myocyte enhancer factor 2 (MEF2) family of MADS-box transcription factors (Puri & Sartorelli, 2000; Taylor & Hughes, 2017). Although many studies have advanced our understanding of myogenesis in both vertebrates and invertebrates, most have focused on differentiation or regeneration in established, adult muscle tissue or have investigated signaling dynamics in isolated, homogeneous systems. As a result, much less is known about how muscle progenitor fate is regulated *in vivo* during the initial specification of satellite cells within a spatially organized, developing niche. In particular, it has remained unclear how signaling gradients in the native tissue environment restrict the activation of key myogenic effectors to maintain subsets of progenitors in a less committed state during early cell fate specification. This regulation serves both to preserve a naïve, undifferentiated population for satellite cell selection and to coordinate myoblast fusion through staggered lineage commitment.

Among these effectors, the MEF2 family plays a conserved and essential role in activating muscle-specific gene expression during differentiation (reviewed in Taylor & Hughes, 2017). While the MEF2 family has been extensively studied in differentiated muscle, it remains unclear how its expression is selectively restricted within progenitor populations in spatially organized developmental niches, particularly during the time when subsets of progenitors are either selected as satellite cells or begin differentiating. Such spatial restriction is likely essential not only to prevent premature or widespread differentiation, thereby preserving a naïve pool for long-term stem cell potential, but also to enable staggered commitment of differentiating progenitors, ensuring proper temporal coordination of myoblast fusion and tissue assembly. To address this question in a spatially tractable *in vivo* system, we investigated the regulation of Mef2 during muscle development in *Drosophila*. We focused on larval muscle progenitors, which give rise to the adult flight muscles in a process that parallels vertebrate skeletal muscle development (Taylor & Hughes, 2017), to examine how expression of *Mef2*, the sole *Drosophila* ortholog of the vertebrate MEF2 family, is regulated during the initial stages of progenitor fate specification. *Drosophila* development proceeds through embryogenesis, three larval instar stages, pupation, and adulthood. The larval muscle progenitors, also referred to as adult muscle progenitors (AMPs), adepithelial cells, or myoblasts, are specified during embryogenesis (Bate et al., 1991) and remain closely associated with an epithelial niche, the wing imaginal disc, which gives rise to the adult wing and associated thoracic structures (Tripathi & Irvine, 2022). This system offers unique spatial and temporal resolution for studying muscle progenitor fate within a native developmental context. During the larval stages, the progenitor pool expands in response to niche-derived signals (Everetts et al., 2021; Gunage et al., 2014; Sudarsan et al., 2001; Vishal et al., 2020). By the third instar larval stage, two distinct subpopulations can be identified: indirect flight muscle progenitors (IFM-MPs) and direct flight muscle progenitors (DFM-MPs). These are distinguished by differential expression of the transcription factor Cut, with IFM-MPs showing low levels and DFM-MPs showing high levels of Cut (Sudarsan et al., 2001) (Fig. 1A, 1A’, 1C’). By early third instar, muscle progenitors form a multilayered tissue structure along the proximal-distal axis of the muscle stem cell niche, as also visualized in the orthogonal view (Fig. 1A’). During pupal stages, these cells either migrate and fuse with persistent scaffolds from the embryonic musculature or fuse to generate the adult flight muscles *de novo*, resulting in the dorsal longitudinal and dorsal ventral indirect flight muscles (DLM- and DVM- IFMs, respectively), as well as the direct flight muscles (DFMs) (Fig. 1B, reviewed in Laurichesse & Soler, 2020).

**Figure 1:**
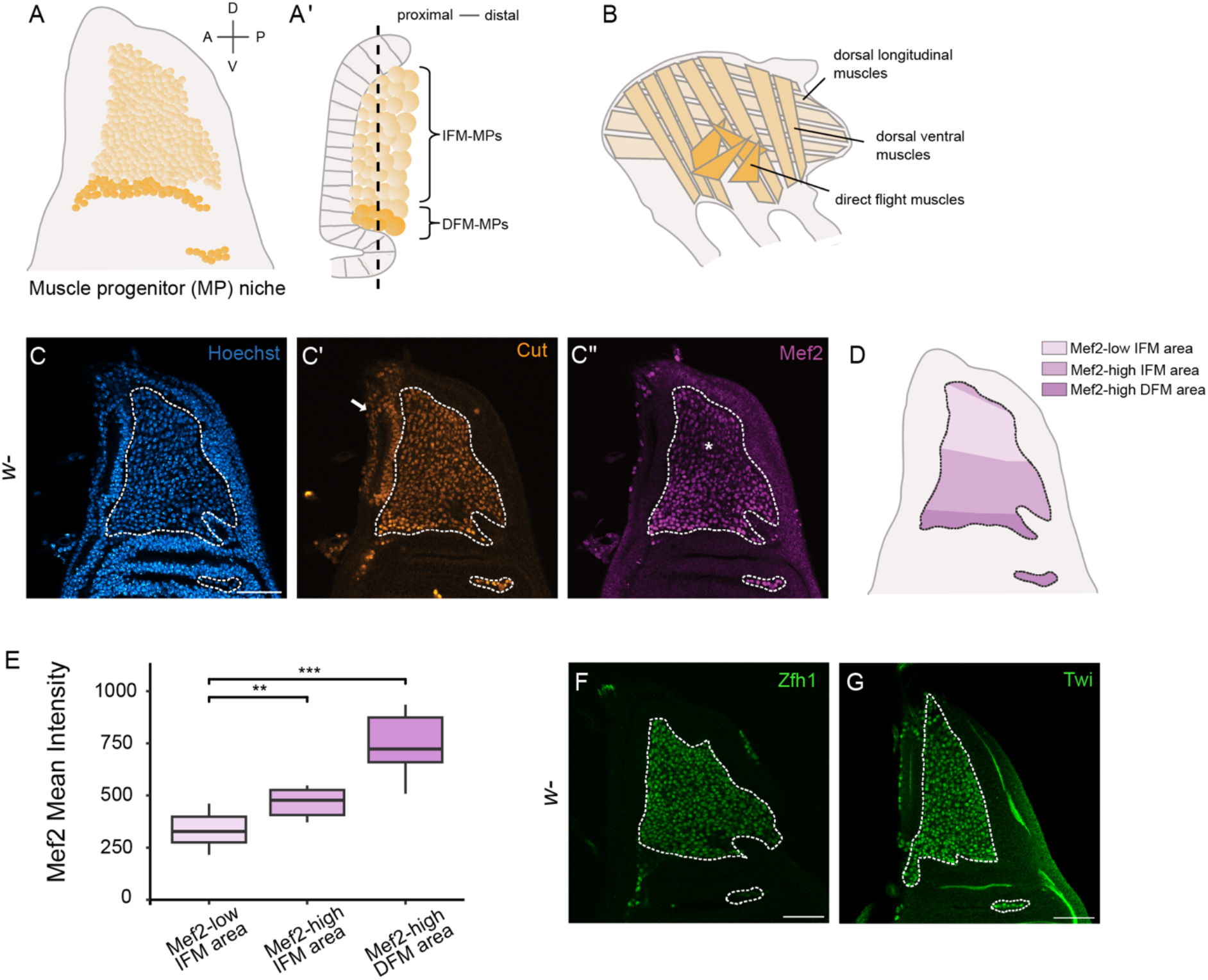
A spatially restricted Mef2-low population exists within the larval muscle progenitor niche of *Drosophila*. **A)** Scheme of the muscle progenitor (MP) niche in late *Drosophila* larvae, showing a single z-plane, and an orthogonal view **(A’)** highlighting the layered structure of the MP pool. The dashed line marks the layer analyzed in this study. Based on Cut levels, IFM-MPs are shown in light orange and DFM-MPs in dark orange. **B)** Schematic of adult flight muscles of *Drosophila*. The dorsal longitudinal (DLM) muscles and dorsal ventral muscles (DVM), which are indirect flight muscles, are derived from IFM-MPs. DFM-MPs give rise to the direct flight muscles (DFM). **C)** Confocal image showing a single z-plane of the larval MP niche. Nuclei are labelled with Hoechst. **C’)** Cut antibody staining marks both IFM- and DFM-MPs, as well as a group of epithelial cells (highlighted with an arrow) that are Cut-positive but not MPs. **C”)** Immunostaining for Mef2 reveals a spatially heterogeneous Mef2 pattern within MPs (n=8). Dorsal IFM-MPs show low Mef2 levels (highlighted with an asterisk), while ventral IFM-MPs and DFM-MPs show higher levels, forming a gradient of Mef2. **D)** Schematic summarizing the spatial organization of Mef2 levels. MPs are divided into three regions based on Mef2 intensity: Mef2-low IFM-MPs (dorsal), Mef2-high IFM-MPs (ventral), and Mef2-high DFM-MPs. **E)** Quantification of nuclear Mef2 intensity across the three regions (n=8), using single-cell nuclear intensity analysis. Wilcoxon statistical test was performed. **F)** Immunostaining for Zfh1 (n=5) and Twist (Twi; n=4) shows no spatial restriction correlating with the Mef2-low IFM-MP region. All confocal images show a single z-plane of the MP niche, and the MP region is outline by a dashed line. Scale bars: 50 µm (confocal images).

We focused our study on the third instar larval stage, a time when Mef2 levels are rising but remain below the threshold for fusion or differentiation (Vishal et al., 2023). *Mef2* expression increases during muscle development, and is detectable in the larval muscle progenitor niche by the third instar larval stage (Cripps et al., 2004; Soler & Taylor, 2009). This upregulation is controlled by a combination of intrinsic and extrinsic cues. One key factor is the basic helix–loop–helix (bHLH) transcription factor Twist, which acts as a repressor of differentiation during early larval development. Upon the ecdysone pulse in late wandering third instar larvae, Twist switches roles and activates *Mef2* expression in the muscle progenitors (Cripps et al., 1998; Lovato et al., 2005). We reasoned that this developmental period provides a unique opportunity to understand how spatial cues restrict Mef2 expression to maintain a less committed state in some progenitors while allowing others to progress towards differentiation.

In line with this, our study reveals a spatially restricted Mef2-low subpopulation of IFM-MPs in the dorsal region of the niche. Using MP-specific genetic perturbations, spatial phenotypic analysis, and single-cell transcriptomics, we show that Wnt/β- catenin signaling regulates *Mef2* transcriptional levels and sustains *zfh1-long* expression across the progenitor pool. This regulatory relationship suggests a role for Wnt/β-catenin signaling in supporting MP maintenance and contributing to heterogeneity in commitment states within the niche. By shaping how many progenitors remain less committed, this mechanism may ultimately influence the number of adult muscle stem cells and impact long-term muscle function and regenerative capacity, a strategy that may be conserved in other systems where Wnt signaling regulates muscle stem cell dynamics.

## Results

To address whether and how *Mef2* expression is spatially restricted within undifferentiated muscle progenitors during early fate specification, we examined Mef2 protein distribution in the larval muscle progenitor (MP) niche at the late third instar stage, a developmental window when Mef2 levels begin to rise but precede overt differentiation. Muscle progenitors were labeled using Cut (Fig. 1C’), which is detected at low level in indirect flight muscle progenitors (IFM-MPs) and at high levels in direct flight muscle progenitors (DFM-MPs). Cut also labels a group of epithelial cells (highlighted by an arrow in Fig. 1C’), which are spatially distinct from the MPs. Immunostaining revealed a previously unnoticed heterogeneous distribution of Mef2 protein within the muscle progenitor pool. In particular, dorsally located IFM-MPs exhibited lower levels of Mef2 than the more ventrally positioned IFM-MPs, while DFM- MPs showed the highest level of Mef2 protein (Fig. 1C”). This results in a gradient of Mef2 level that decreases towards the dorsal region, suggesting spatial regulation of differentiation competence. For quantification, we defined three regions within the MP pool based on Mef2 intensity: Mef2-low IFM-MP, Mef2-high IFM-MPs and Mef2-high DFM-MPs (Fig. 1D). Measurement of nuclear Mef2 intensity confirmed that Mef2 levels are significantly different between these regions (Fig. 1E). Since Mef2 levels generally rise over developmental time (Cripps et al., 2004; Soler & Taylor, 2009), we hypothesized that the ventral and DFM-MPs likely reflect the default progression toward differentiation. In contrast, the persistence of Mef2-low levels in a spatially defined dorsal subset suggests that active repression, rather than localized activation, underlies the observed heterogeneity.

To assess whether known transcriptional repressors of Mef2 could account for its spatial pattern, we stained for Zfh1 and Twist, both of which have been proposed to antagonize Mef2 expression (Postigo et al., 1999). However, neither Zfh1 (Fig. 1F) nor Twist (Fig. 1G) displayed higher expression in the Mef2-low IFM-MP region, indicating that these factors alone are unlikely to underlie the regional Mef2 heterogeneity observed in this study. Notably, occasional Mef2-low cells were also observed outside the dorsal domain, in both Mef2-high IFM- and DFM-MPs, particularly among phospho-Histone H3 (PH3)-positive mitotic cells (Supp. Fig1 A-C). This finding is consistent with the known degradation of MEF2C, one of the four MEF2 family members in vertebrates, during mitosis (Badodi et al., 2015). However, PH3 staining did not reveal any obvious spatial differences in mitotic cell distributions, suggesting that mitotic degradation alone does not account for the spatially clustered Mef2-low population (Supp. Fig1 A-C). This contrasts with the consistently observed, spatially organized Mef2-low domain analyzed here.

Notably, the spatially clustered Mef2-low domain resembles a subset of IFM- MPs reported by Boukhatmi and Bray (Boukhatmi & Bray, 2018), in which a specific enhancer of the *zfh1* gene, termed “Enh3”, is active. Although this enhancer does not fully reflect the expression of either *the zfh1-long* or *zfh1-short* isoform during larval stages, it nevertheless marks a subset of progenitor cells that were shown by Boukhatmi and Bray to remain unfused during pupal stages and contribute to the pool of satellite-like cells in the adult, which are characterized by low *Mef2* expression and continued Zfh1 level. (Boukhatmi & Bray, 2018).

To test whether the Mef2-low population corresponds to this Enh3+ domain, we used an Enh3-Gal4 driver to express nuclear GFP and co-stained the larval muscle progenitors for Mef2 protein. Confocal imaging confirmed that the Enh3-GFP signal overlapped spatially with the region of reduced Mef2 level (Fig. 2). This suggests that Enh3 is active within the Mef2-low domain, highlighting a potential early mechanism by which differentiation is selectively delayed or prevented in a subset of muscle progenitors. Such regulation may contribute both to the staggered timing of myoblast fusion and to the maintenance of satellite-like cells into adulthood.

**Figure 2:**
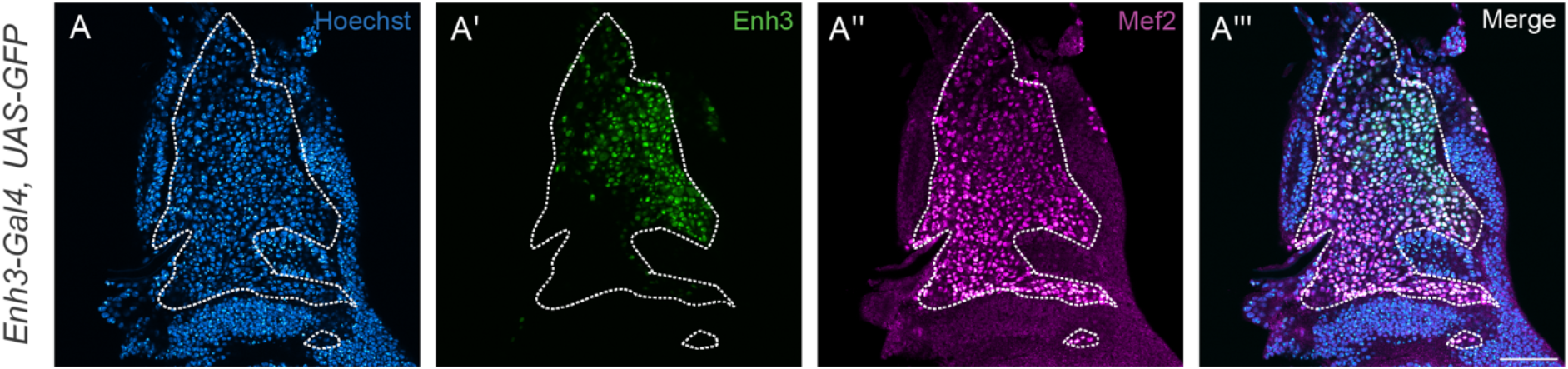
Enh3 activity overlaps with the Mef2-low IFM-MP subset, previously linked to adult satellite-like cells through Enh3 lineage. Confocal single-plane image of the larval muscle progenitor (MP) niche A’) Hoechst labels all nuclei, **A”)** GFP fluorescence driven by Enh3-Gal4 (*zfh1* Enh3 element) is observed in a spatially restricted region of the MP niche. **A”’)** Mef2 antibody staining shows reduced levels within the Enh3-positive domain. **A””)** Merged image shows the overlap between Enh3- driven GFP expression and the Mef2-low IFM-MP subset (n=5). The MP region is outlined by a dashed line. Scale bar: 50 µm.

Notch signaling plays a central role in maintaining progenitor identity, in part through sustaining Zfh1 level (Boukhatmi & Bray, 2018). Given this role, and the presence of a binding site for Suppressor of Hairless [Su(H)], the transcriptional effector of the Notch signaling pathway, in the Enh3 enhancer, we next asked whether Notch signaling contributes to the spatial regulation of Mef2 within the muscle progenitors. To test this, we used the MP-specific 1151-Gal4 driver (Anant et al., 1998) to express RNAi against the *Notch* receptor, thereby blocking pathway activation. However, perturbing Notch signaling in this way did not disrupt the spatial restriction of the Mef2-low domain (Suppl. Fig. 2A, B). This suggests that additional, spatially localized cues are required to establish regional differences in differentiation competence.

To determine whether the observed heterogeneity in Mef2 protein levels is accompanied by differences in *Mef2* transcription, we performed single-molecule FISH (smFISH) against *Mef2* mRNA in the third instar larval muscle progenitor niche, confirming that the spatial Mef2 pattern reflects transcriptional differences (Fig. 3A). This, in turn, enabled us to use available single-cell transcriptome data to investigate the regulatory logic underlying Mef2 heterogeneity. To do this, we integrated and re- analyzed published single-cell RNA-seq (scRNA-seq) datasets acquired using 10x Genomics platform from Bageritz et al. (Bageritz et al., 2019) and Everetts et al. (Everetts et al., 2021). The integrated UMAP representation (Fig. 3B) shows the distribution of identified clusters across the combined dataset after cell cycle regression. Replicate agreement was strong, with comparable cluster proportions and gene and transcript complexity across samples (Suppl. Fig. 3A-C), indicating successful integration and robust clustering. Our analysis focused on identifying and characterizing the Mef2-low population. While our clustering resolution differs from the higher granularity reported by Zappia et al. (2023), who annotated five IFM subclusters in their Drop-seq-generated scRNA-seq data, we observed a correspondence between our Mef2-low clusters and their identified IFM-3 and IFM-4 clusters (Suppl. Fig. 3D). Importantly, the high transcriptome coverage in our combined dataset, with an average of approximately 2,500 genes and 16,000 transcripts detected per cell, enabled the detection of the more weakly expressed transcripts, such as *Mef2*. Indeed, in all three biological replicates, clusters 0 and 3 consistently exhibited the lowest *Mef2* transcript levels (Fig. 3C). Although regression of cell cycle effects was generally effective, clusters 0 and 3, which were similar in transcript counts (Suppl. Fig. 3E), showed differences in S and G2/M scores (Suppl. Fig. 3F). This indicates that residual cell cycle effects may persist despite correction with the current gene set. Nevertheless, the integrated dataset provides a robust framework for further analysis of the regulatory inputs defining the Mef2-low population. Among the top markers enriched in the Mef2-low IFM-MP population compared to Mef2-high IFM-MPs, we identified *rotund* (*rn*), which was highly specific to this group (Fig. 3D; Suppl. Fig. 3G). We confirmed the spatial overlap in gene expression pattern between *rn* and Mef2- low cells using a rn-Gal4 > UAS-RFP reporter (Suppl. Fig. 3H), which showed dorsal MP expression pattern consistent with previously published data (pre-print from Everetts et al., 2020; St Pierre et al., 2002). This validation confirms that the spatially restricted Mef2-low population is preserved in our scRNAs-seq data, linking transcriptomic profiles to anatomically defined subsets of IFM-MPs.

**Figure 3:**
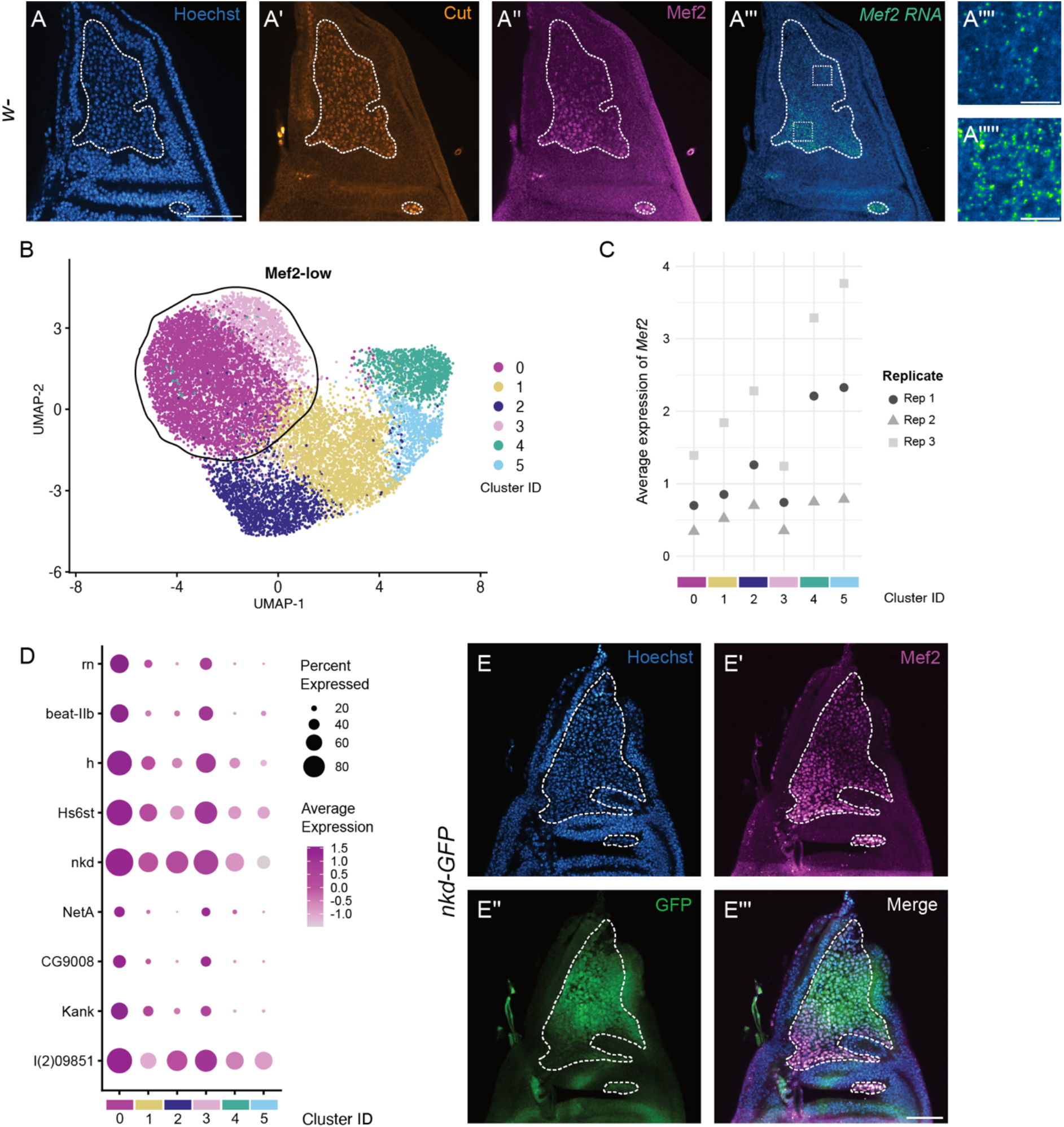
Mef2 heterogeneity is regulated at the transcriptional level in *Drosophila* larval muscle progenitors. **A)** Confocal single-plane image of the muscle progenitor niche (MP) in wandering third instar (L3) larvae showing antibody staining for Cut **(A’)** and Mef2 **(A”)** proteins, alongside smFISH detection of *Mef2* mRNA **(A”’)** with close-ups of the probe in the Mef2-low **(A””)** and Mef2-high IFM- MPs **(A””’)**. The MP region is outlined by a dashed line. Scale bar MP niche: 50 µm, scale bar close-ups: 5 µm (n= 2). **B)** UMAP plot of integrated single-cell RNA-sequencing data sets from wildtype wandering L3 larvae. Mef2-low MPs are found in two distinct clusters. **C)** Average *Mef2* expression per cluster across three biological replicates. Clusters 0 and 3 exhibit the lowest *Mef2* expression levels. **D)** Dot plot showing the top marker genes for the Mef2-low clusters (cluster 0 and 3) compared to remaining IFM-MP clusters (cluster 1 and 2). Marker genes have an average log2 fold change ≥ 0.7 and an adjusted p-value ≤ 0.1 in all three replicates. **E)** Confocal single-plane image of Mef2 immunostaining in a nkd-GFP reporter line. GFP expression overlaps with the Mef2-low IFM-MP region of the muscle progenitor niche (n=5). The MP region is outlined by a dashed line. Scale bar: 50 µm.

We next asked whether a signaling pathway with regionally enriched activity might underlie the localized *Mef2* repression observed in the dorsal IFM-MPs. Among the most enriched genes in the Mef2-low population was the Wnt/β-catenin target *naked cuticle* (*nkd*) (Zeng et al., 2000; Fig. 3D), whose expression level inversely correlated with *Mef2*. We validated the graded *nkd* expression using a nkd- GFP reporter, which showed high activity in the dorsal IFM-MP region corresponding to Mef2-low IFM-MP cells (Fig. 3E), and lower activity in the Mef2-high IFM-MP population. This anti-correlation suggested a potential role for Wnt signaling in regulating Mef2 heterogeneity. Wnt/β-catenin signaling is initiated when Wnt ligands bind to Frizzled (Fz) receptors and the co-receptor Arrow (the *Drosophila* homolog of vertebrate LRP5/6). This interaction leads to stabilization of the transcriptional co- activator armadillo (arm, *Drosophila* β-catenin), which translocate into the nucleus and partners with pangolin (pan, *Drosophila* TCF) to activate target gene expression (Fig. 4A). While this pathway is classically associated with transcriptional activation, the observed anti-correlation between *nkd* and *Mef2* expression suggests that Wnt signaling may directly or indirectly repress Mef2, for example by activating an intermediate repressor. To test whether Wnt signaling contributes to the regulation of Mef2 levels in muscle progenitors, we used the MP-specific 1151-Gal4 driver to express *arm* RNAi and inhibit pathway activity. This resulted in a significant increase in Mef2 protein levels, most strongly in the previously defined Mef2-low dorsal region, leading to an overall homogenization of Mef2 level across the MP pool (Fig. 4B–C; quantified in Fig. 4E). Knockdown efficiency was quantified, revealing a reduction of Arm protein to approximately 45% following RNAi-mediated *arm* knockdown compared to the control (Fig. 4D).

**Figure 4:**
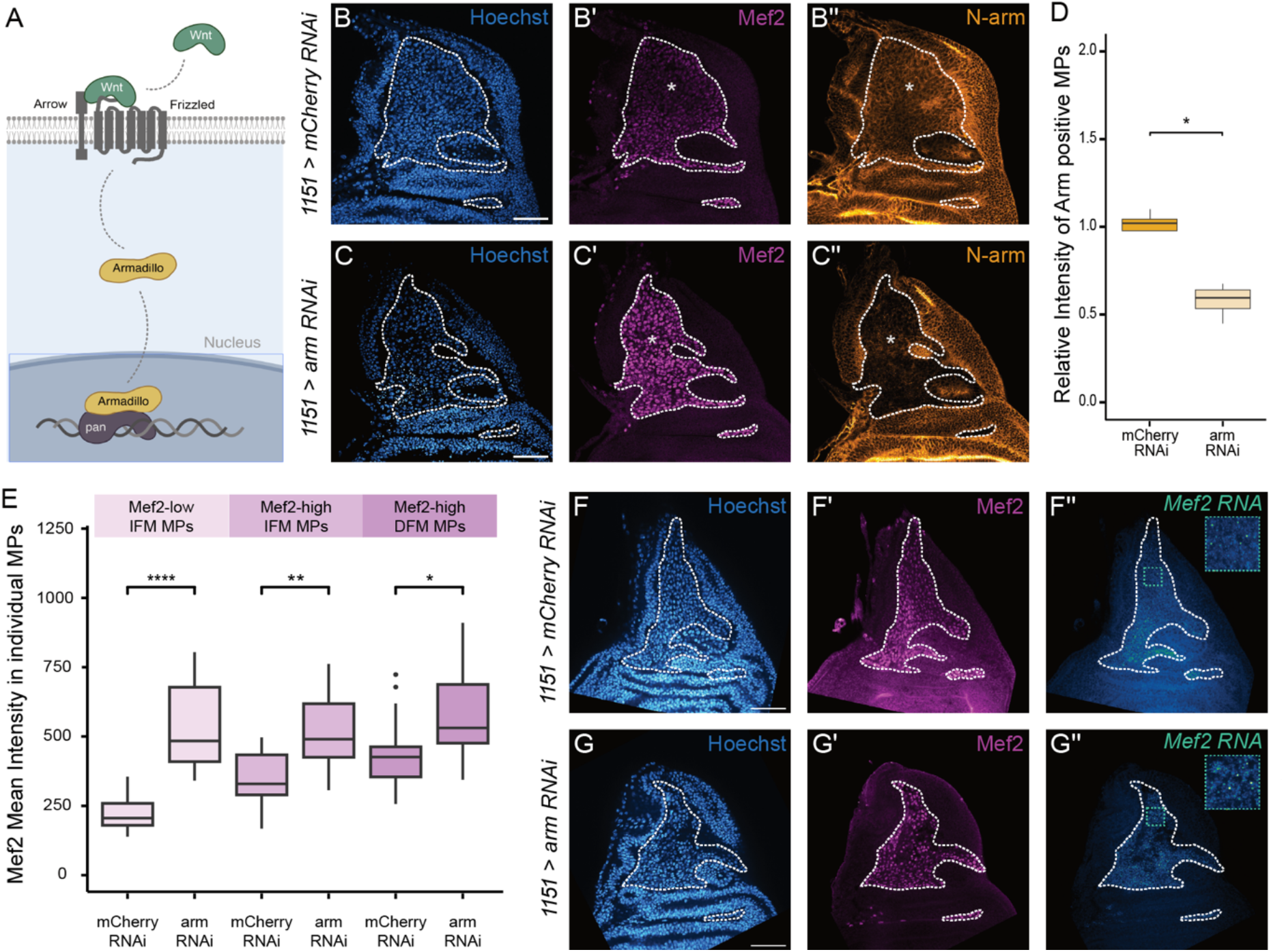
Wnt/β-catenin signaling regulates Mef2 transcription in *Drosophila* muscle progenitors. **A)** Simplified scheme of the Armadillo (Arm, *Drosophila* β-catenin)-dependent Wnt signaling pathway. Binding of Wnt ligands to Frizzled (Fz) receptors and the co-receptor Arrow leads to stabilization and nuclear translocation of Arm, which partners with Pangolin (Pan; *Drosophila* TCF) to activate transcriptional targets. **B-C)** Immunofluorescence staining of Mef2 protein after MP-specific *arm* knockdown using 1151-Gal4 driver. Mef2 levels are increased, particularly in the dorsal Mef2-low region, but also across broader areas of the IFM-MP pool, leading to a more uniform Mef2 pattern. Representative single-plane images are shown, the MP region outlined by a dashed line. The Mef2-low area is marked by an asterisk. Scale bar: 50 µm. **D)** Quantification of *armadillo* knockdown efficiency based on Arm protein intensity in the MP area. Protein levels are significantly reduced to ∼45% relative to control (n=4). Wilcoxon statistical test was performed. **E)** Quantification of Mef2 protein level after *armadillo* knockdown in the three defined MP regions (Mef2-low IFM-MPs, Mef2-high IFM-MPs, and Mef2-high DFM-MPs), using single-cell nuclear intensity analysis. Mef2 levels are significantly elevated in all regions following *arm* knockdown (control: n=13, RNAi: n=12, across two independent experiments). Images and quantification are from independent biological replicates. Statistical significance was assessed using the Wilcoxon test. **F-G)** Representative single-plane images showing Mef2 protein (immunofluorescence) and *Mef2* mRNA (smFISH) following *arm* knockdown (control: n=9; *arm* knockdown: n=6, across two independent experiments). *Mef2* transcript levels are increased. The MP region is outlined by a dashed line and close-ups of the Mef2-low IFM-MPs is shown in the right corner. Scale bar: 50 µm.

To assess whether the observed phenotype results from impaired Wnt/β- catenin signaling rather than disrupted cell–cell adhesion, due to Armadillo’s additional role at the membrane (Cox et al., 1996), we expressed a dominant-negative form of TCF (TCF-DN) using the 1151-Gal4 driver. This recapitulated the phenotype (Suppl. Fig.4 A-C), indicating that the effect is mediated via Wnt/β-catenin signaling. Consistent with this, smFISH analysis revealed that *nkd* expression was significantly reduced upon *arm* knockdown (Suppl. Fig. 4 D, E), further supporting disruption of the Arm–TCF signaling axis. Given that Mef2 heterogeneity is present at the transcript level (Fig. 3A), we next asked whether Wnt/β-catenin signaling regulates *Mef2* at the transcriptional level. To address this, we performed smFISH against *Mef2* mRNA in control (mCherry) and *arm* RNAi condition. *Mef2* transcript levels were elevated upon *arm* knockdown (Fig. 4F, G), demonstrating that Wnt/β-catenin signaling directly or indirectly represses *Mef2* transcription, with the strongest effect observed in regions of high Wnt activity as inferred from *nkd* expression pattern (see Fig. 3E). To test whether Wnt/β-catenin signaling represses Mef2 through known antagonists, we assessed Twist and Zfh1 protein levels following *arm* knockdown. Neither factor showed detectable changes in the Mef2-low area linked to protein abundance or spatial distribution (Suppl. Fig. 5A–D), indicating that Wnt/β-catenin signaling does not regulate these factors at the protein level at the larval stage. Since neither is downregulated upon *arm* knockdown, they are unlikely to mediate the increased *Mef2* expression observed in this context. While Notch signaling regulates *zfh1-short* expression during the pupal stage (Boukhatmi & Bray, 2018), the upstream signals that regulate *zfh1* transcription during larval stages remain unclear. To address whether Wnt/β-catenin signaling contributes to early *zfh1* regulation, and by extension to the selection and maintenance of satellite-like cells, we next asked whether Wnt/β- catenin activity affects *zfh1* expression. Although Zfh1 protein levels remain uniform during larval stages and do not change upon *arm* RNAi (Suppl. Fig. 5C, D), we considered the possibility that Wnt signaling may regulate *zfh1* at the transcriptional level, with effects not yet apparent at the protein level. Such early transcriptional input could help stabilize *zfh1* expression in anticipation of the pupal transition, when Zfh1 degradation begins in differentiating cells. In this context, Wnt signaling may act in parallel to or independently of Notch to prime satellite-like cells for future maintenance. To test whether canonical Wnt signaling influences *zfh1* transcription, we performed smFISH against the *zfh1-long* isoform in late wandering third instar larvae. Upon *arm* knockdown using the 1151-Gal4 driver, *zfh1-long* transcript levels were visibly reduced compared to controls (Fig. 5A, B). Quantification based on smFISH puncta counting revealed an approximate 50% reduction in transcript abundance (Fig. 5C), indicating that Wnt /β-catenin signaling is required to maintain *zfh1-long* transcription during this stage. Attempts to quantify *zfh1-short* mRNA using isoform-specific probes were inconclusive due to limited probe specificity. The observed reduction in *zfh1-long* mRNA is likely to compromise Zfh1 protein stability during pupal stages, potentially destabilizing the undifferentiated state.

**Figure 5:**
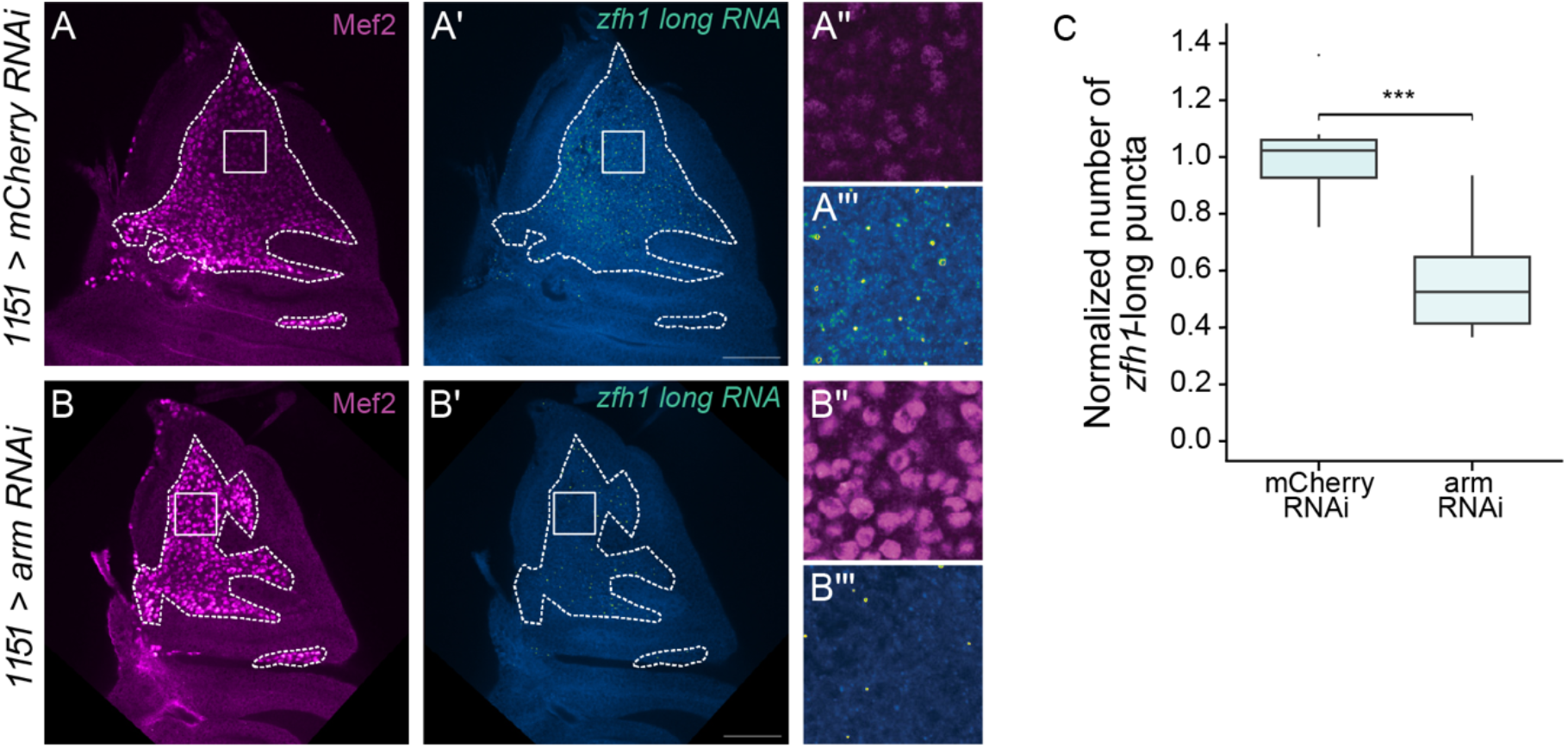
Wnt/β-catenin signaling maintains *zfh1-long* transcription in larval muscle progenitors of *Drosophila*. **A-B)** Representative single-plane confocal images of the larval muscle progenitor niche (MP) in control **(A)** and *arm* knockdown **(B)** conditions using the MP-specific 1151-Gal4 driver. Mef2 protein is detected by immunofluorescence **(A and B)**, and and *zfh1-long* transcripts are visualized by smFISH **(A’, B’)**. Close-ups of the dorsal Mef2-low region in control **(A” and A”’)** and knockdown **(B” and B”’)** samples, respectively, show a clear reduction in *zfh1-long* transcript level after *arm* knockdown, while Mef2 protein levels are elevated. The MP region is outlined by a dashed line. Scale bar: 50 µm. **C)** Quantification of smFISH signal shows a significant reduction in *zfh1-long* transcript abundance following MP-specific (1151-Gal4 driver) *arm* knockdown (n=10, across two independent experiments). Wilcoxon statistical test was performed.

Together with our finding that Wnt/β-catenin signaling represses *Mef2* transcription, these data support a model in which Wnt signaling contributes to muscle progenitor maintenance through dual regulation of *zfh1* and *Mef2*. In particular, the spatially restricted Wnt-high domain exhibits strongest *Mef2* repression, consistent with a role in preserving a naïve, undifferentiated subpopulation for satellite-like cell selection. In line with this idea, overexpression of *Mef2* in the Mef2-low region using rn-Gal4 did not alter the number of DLM or DVM fibers, as assessed by micro-CT (Suppl. Fig. 6A, B), suggesting that fusion capacity remains intact. However, interpretation is limited by the small sample size due to impaired eclosion, and future studies with larger cohorts and direct assessment of satellite-like cell number, especially during the pupal transition, will be important to clarify the functional significance of this progenitor subpopulation.

## Discussion

Despite significant advances in understanding myogenic differentiation and adult stem cell maintenance, the mechanisms that coordinate early lineage commitment with the preservation of undifferentiated progenitors in a spatially organized developmental niche remain incompletely understood. In particular, the process that restrict the activity of key myogenic effectors such as Mef2 during the initial selection of future satellite-like cells are not well characterized *in vivo*. Our study addresses this gap by demonstrating that graded Wnt/β-catenin signaling activity modulates *Mef2* transcription levels and sustains *zfh1* expression across *Drosophila* larval muscle progenitor pool, thereby establishing spatial heterogeneity in differentiation competence.

The spatial co-occurrence of high Wnt signaling, low *Mef2* expression, and activity of the *zfh1* enhancer (Enh3) in a subpopulation of indirect flight muscle progenitors (IFM-MPs), with Enh3 activity shown by lineage analysis to give rise to satellite-like cells, suggests that Wnt signaling plays an early instructive role in initiating the selection of the satellite-like cells. This Wnt-mediated mechanism may act upstream of, or in concert with, Notch signaling during pupal development, with both pathways contributing to the establishment of the satellite-like cell pool required for adult tissue regeneration.

### Low Mef2 level as an early mechanism for satellite-like cell selection

During *Drosophila* development, two waves of myogenesis occur. The first wave in the embryo produces larval muscles, while a subset of myogenic cells (muscle progenitors, MPs) is specified but prevented from differentiating by the transcription factors Twist and Notch. These progenitors remain undifferentiated, proliferate during the larval stages, and contribute to the second wave of myogenesis during pupal stages, when the adult musculature is formed (reviewed in Laurichesse & Soler, 2020). Until recently, it was widely assumed that all muscle progenitors ultimately differentiate and fuse to form multinucleated fibers. However, this view was challenged by the discovery of a distinct subpopulation of satellite-like cells in *Drosophila* by Chaturvedi et al. (Chaturvedi et al., 2017). These adult stem cells are characterized by persistent expression of the stemness factor *zfh1*. During pupal development, Notch signaling plays a key role in maintaining *zfh1* expression, thereby protecting this subset of MPs from differentiation. Consequently, satellite-like cells retain high Zfh1 protein and exhibit low levels of Mef2, a key transcriptional activator of muscle differentiation. Building on the observation that satellite-like cells are characterized by low Mef2 levels, we identified a larval MP subpopulation among the IFM-MPs in the dorsal muscle progenitor niche that exhibits low *Mef2* expression and, together with neighboring MPs, high Zfh1 protein levels, indicating that these cells are held back from lineage progression in anticipation of future adult stem cell function.

While earlier work by Gunthorpe and colleagues (Gunthorpe et al., 1999) demonstrated that varying levels of Mef2 activity contribute to the specification of different somatic muscles, raising the possibility that the observed Mef2-low population among the IFM-MPs could represent an early separation of DLM (dorsolongitudinal) and DVM (dorsoventral) IFM fiber lineages, our *Mef2* overexpression experiments argue against this. Specifically, forcing Mef2 expression in the Mef2-low MPs did not seem to alter the number of either fiber type, suggesting that the Mef2 levels in these cells are unlikely to drive DLM or DVM lineage specification. Instead, the Mef2-low domain overlaps with activity of the *zfh1* Enh3 enhancer, previously shown by genetic lineage analysis to mark progenitors that give rise to adult satellite-like cells (Boukhatmi & Bray, 2018). The spatial convergence of low *Mef2* expression, high Zfh1 protein levels, and Enh3 activity supports the idea that these cells represent early-selected satellite-like progenitors.

At first glance, the presence of distinct Mef2 expression levels among larval IFM-MPs may seem unexpected, given that all muscle progenitors at this stage are undifferentiated, not expressing differentiation markers such as MHC (Vishal et al., 2023). However, we propose that these differences in Mef2 levels reflect a hierarchical regulation within a pre-differentiation state: transcriptional repression provides a spatially stable and robust foundation for progenitor identity, particularly in the Mef2- low domain, while post-translational mechanisms, such as phosphorylation (Vishal et al., 2023), could represent a more transient and readily reversible layer of regulation in other regions of the tissue. These observations are consistent with a model in which transcriptional repression establishes a robust progenitor state in the Mef2-low domain, while more dynamic post-translational mechanisms such as phosphorylation could help maint other cells in a pre-differentiation state below the threshold required for full commitment. The observed spatial heterogeneity in *Mef2* expression among larval IFM-MPs may point to early regulatory differences that influence progenitor potency or protective states, although the lineage relationships among IFM-MPs are not yet fully understood. Zappia et al. started to address this question from a transcriptomic perspective by applying pseudotime analysis to single-cell RNA-seq data, inferring a developmental trajectory based on transcriptional similarity. In their analysis, the Mef2-low population (designated “IFM-4”) occupies an intermediate position along this pseudotemporal axis, suggesting a transition state within the developmental trajectory. Whether *Drosophila* satellite-like cells emerge from intermediate myogenic states or are protected from commitment remains unclear. Spatially and temporally controlled genetic lineage analysis will be key to clarifying their developmental origin.

### Wnt/β-catenin signaling as a key regulator of Mef2 and zfh1 transcription

To investigate the molecular mechanism underlying spatial *Mef2* repression, we made us of our integrated single-cell RNA-seq datasets allowing us to identify two transcriptionally distinct Mef2-low clusters. Among the top markers of the Mef2-low cells were *rotund* (*rn*) and *hairy* (*h*), both of which have known spatial expression patterns within the IFMs (pre-print of Everetts et al., 2020; St Pierre et al., 2002), reinforcing the link between these transcriptional profiles and the dorsal IFM-MP domain *in vivo*. This enabled us to investigate localized regulatory mechanisms impacting the dorsal IFM-MP domain. Since *hairy* is a classical Notch target gene and the *zfh1* Enh3 enhancer active in the Mef2-low region was shown to be Notch-responsive (Boukhatmi & Bray, 2018), we considered whether Notch signaling might mediate localized Mef2 repression during development. Notch knockdown did not disrupt the restriction of the Mef2-low domain, the spatial heterogeneity remains intact with this domain still clearly detectable. This suggests that, although Notch signaling may contribute to aspect of progenitor regulation (Gunage et al., 2014), it is not required for establishing or maintaining the spatial pattern of Mef2 at the larval stage. Since we did not directly assess *hairy* expression in this context, it remains possible that *hairy* expression is regulated independently of Notch during this developmental window.

Notch signaling may instead play a more prominent role later in satellite-like cell selection during pupal stages, as previously proposed by Boukhatmi and Bray (Boukhatmi & Bray, 2018). The early activity of the Enh3 enhancer in Mef2-low cells thus suggest that the satellite-like progenitor identity is established before the onset of Notch signaling. In fact, the known spatial expression of *Wingless* (*Wg*) in the adjacent epithelial niche close to the dorsal Mef2-low IFM-MPs in earlier stages (Mirth et al., 2009), and our data provide compelling evidence that Wnt/β-catenin signaling plays a central role in repressing *Mef2* expression. We observed dorsal enrichment of two Wnt-associated genes: *naked cuticle* (*nkd*), a well-established Wnt/β-catenin target (Zeng et al., 2000), and *Hs6st*, a heparan sulfate 6-O-sulfotransferase known to enhance Wnt ligand–receptor interactions by modifying co-receptors (Dani et al., 2012). Importantly, inhibition of Wnt signaling specifically in the MPs resulted in elevated *Mef2* transcript and protein levels particular in the Mef2-low MPs thereby abolishing the spatial Mef2 patterning. These manipulations also reduced *nkd* expression, confirming effective Wnt pathway inhibition. Together, these findings establish Wnt/β-catenin signaling as a transcriptional repressor of *Mef2 in vivo*.

Importantly the localized high activity of Wnt pathway in the dorsal IFM-MP domain leads to a pattern of heterogeneous Mef2 level. Mechanistically, Wnt signaling may repress *Mef2* through activity of a transcriptional repressor. Although Twist and Zfh1 protein levels remain unchanged upon Wnt pathway inhibition, their activity could be regulated via post-translational modifications, including phosphorylation, as suggested for Twist (Balakrishnan et al., 2021). In addition, the activity of transcriptional repressors like Twist is often context-dependent and can rely on interactions with specific cofactors. Notably, Twist requires heterodimerization with the bHLH protein Daughterless to function as a repressor (Castanon et al., 2001). This raises the possibility that Wnt signaling may influence *Mef2* repression not only through transcription factor abundance, but also by modulating the expression, availability, or interaction dynamics of essential cofactors. A more direct mechanism could involve Tcf-mediated transcriptional repression of *Mef2*, as demonstrated in other developmental contexts (Blauwkamp et al., 2008). Alternatively, Wnt signaling might affect *Mef2* expression by modulating the stability or abundance of *Mef2* transcripts. Interestingly, a similar Wnt-mediated repression of Mef2 transcription was reported in mice, where Wnt16 acts to suppress Mef2C in postnatal bone homeostasis (Sebastian et al., 2018). Future work will be needed to identify the precise effectors downstream of Armadillo(β-catenin)/TCF that mediate *Mef2* repression, and to determine whether similar mechanisms operate across species.

In addition to repressing *Mef2*, our data show that Wnt/β-catenin signaling supports transcription of the *zfh1-long* isoform across the IFM-MP population. smFISH analysis revealed widespread *zfh1-long* expression in IFM-MPs, which was reduced upon *armadillo* knockdown, indicating that Wnt signaling promotes its expression. Although we could not reliably quantify *zfh1-short* by smFISH due to limited probe specificity, it remains possible that Wnt signaling also contributes to its regulation during larval stages. This may complement Notch-mediated *zfh1-short* transcriptional regulation observed during the pupal stage (Boukhatmi & Bray, 2018), or reflect a temporally layered regulatory mechanism. In contrast to this Wnt-mediated repression of *Mef2* during early stages, previous work has shown that ecdysone, in cooperation with Twist, promotes an increase in *Mef2* expression at the end of larval development (Lovato et al., 2005), thereby initiating differentiation. Although not shown here, our single-cell data indicate that the ecdysone receptor (EcR) is expressed at lower levels in the Mef2-low population, suggesting reduced sensitivity to ecdysone signaling.

These findings raise the possibility that the Mef2-low subset of IFM-MPs is selectively spared from this differentiation-promoting cue. Previous studies have linked Wg/Wnt signaling to the regulation of the MPs. Sudarsan et al. (Sudarsan et al., 2001) found that Wg mutants have reduced numbers of MPs, which could be caused by altered proliferation as suggested by Gunage et al. (Gunage et al., 2014). In contrast, our data reveal a regional function for Wnt/β-catenin signaling in the dorsal IFM-MP region, where high pathway activity represses *Mef2*, a process suggested to maintain progenitor identity. The observed decrease in MP number following Wnt pathway inhibition may therefore reflect a failure to maintain the undifferentiated progenitor pool, a mechanism that by altering division mode (symmetric vs. asymmetric) and proliferation rate could also affect total MP number. Notably, these findings may reflect a conserved developmental strategy across species. In vertebrates, Hutcheson and colleagues (Hutcheson et al., 2009) demonstrated that β-catenin is essential for maintaining the fetal muscle progenitor pool in developing limbs, likely by modulating differentiation of Pax7+ cells. This parallel suggests that spatially graded Wnt signaling activity may be a broadly utilized mechanism to balance progenitor maintenance and differentiation across species. In light of this, future studies should aim to dissect how region-specific differences in Wnt signaling levels contribute to distinct outcomes, such as stemness maintenance versus lineage commitment, within the *Drosophila* muscle progenitor pool.

## Material and Methods

### Fly maintenance and fly lines

Flies were kept at 25°C in a 12h light-dark cycle. Egg laying was performed for 24h at 25°C and crosses were performed at either 25°C (dnTCF cross, *Mef2* overexpression) or at 29°C (*arm* knockdown, *Notch* knockdown). To investigate the *Mef2* spatial expression on protein and RNA level within MPs in wild type condition we used the *Drosophila* strain w1118 (Fig. 1 and Fig. 3). Cell-type specific gene knockdowns and overexpressions were achieved using the following Gal4-driver and RNAi lines: 1151-Gal4 (Anant et al., 1998), rn-Gal/TM6B (provide by Aurelio Teleman), UAS-mCherry RNAi (BL 35787), UAS-RFP (BL 30556), UAS-arm RNAi (BL 31305), UAS-Notch RNAi (VDRC 100002), UAS-dnTCF (BL 4784), UAS-Mef2 (from Hanh Nguyen). To proof Enh3-Gal4 and *nkd*-GFP reporter line activity, following GFP lines were used: Enh3-Gal4, UAS-GFP line (provided by Hadi Boukhatmi (Boukhatmi & Bray, 2018), *nkd*-GFP reporter line (BL 30664).

### Generation of labelled oligonucleotides for smFISH

The Stellaris™ probe designer (https://www.biosearchtech.com/support/tools/design-software/stellaris-probe-designer) was used to design 18-22 nucleotide long probes targeting *Mef2, zfh1-long* isoform and *nkd* RNA. The specificity of the probes was checked by blasting individual oligonucleotides. Oligonucleotides that detected an off-target gene with more than four oligonucleotides were discarded to prevent the detection of non-specific signals. The final probe set consisted of 40 to 44 specific probes, which were ordered from Eurofins Scientific pre-diluted to 200 pmol/µl. Labelled oligonucleotides for single molecule fluorescence in situ hybridization (smFISH) were custom-made following published protocols (Gáspár et al., 2017, Gáspár et al., 2018), using an Atto-633 as a fluorophore. Specifically, probes were labelled by conjugating Amino-11-ddUTP with the Atto-633 NHS-ester (Sigma-Aldrich), followed by a terminal deoxynucleotidyl transferase reaction. The final probes were purified by ethanol precipitation and resuspended in a total volume of 20 µl water.

### Dissection, immunostaining and in-situ hybridization

All steps for immunostainings were carried out in the dark and with gentle rotation. Wandering third-instar (L3) larvae were dissected in 1x phosphate-buffered saline (PBS) and fixed with 4% paraformaldehyde (PFA, diluted in PBS) for 25 min. Samples were washed three times with PBS containing 0.3% Triton X-100 (PBST) for 10 min each, then blocked with 1% normal goat serum (diluted in PBS) for 1 hour. Primary antibodies were diluted in PBST and incubated with the samples overnight at 4°C. Following incubation, samples were rinsed and washed three times with PBST. Secondary antibodies (Invitrogen, all secondary antibodies diluted 1:500 in PBST, excepted donkey anti-chicken AF488 (1:4000) and Hoechst (Invitrogen, diluted 1:3000 in PBST) were added and incubated for 2-3 hours at room temperature. To prevent cross-reactivity between secondary antibodies during immunostaining combination of Mef2, Cut and PH3, highly cross-absorbed secondary antibodies were used and a sequential staining was carried out.

For immunostaining, the following primary antibodies were used: Rabbit anti- Mef2 (generated by H. T. Nguyen, provided by Katrin Domsch, Heidelberg University, 1:1000), mouse anti-Cut 2B10 (DSHB, 0.4 µg/ml), mouse anti-Armadillo N2 7A1 (DSHB, 0.52 µg/ml), rabbit anti-Zfh1 (gift from Ruth Lehmann, MIT, Cambridge, MA, 1:5000), rabbit anti-Twist (Furlong Lab, EMBL, Heidelberg, 1:1000), Rat anti-PH3 (clone HTA28, abcam, 1:10,000).

To assess transcript levels of *Mef2, nkd*, and *zfh1-long*, smFISH staining was performed. Wandering third-instar larvae were dissected in 1x PBS and fixed for 25 min in 4% PFA. Samples were then washed three times for 10 min each with 0.1% PBST (PBS with 0.1% Triton-X-100), followed by fixation in 70% ethanol for 24 hours at 4°C. For hybridization, fixed samples were incubated for 10 min in 1000 μl Wash buffer A (10% formamide, 2x SSC, 0.1%Triton X-100 in molecular-grade water), and then hybridized overnight at 37°C with shaking (200 rpm), protected from light. Hybridization was carried out in 100-250 μl hybridization solution (10 % formamide, 2x SSC, 10% dextran sulphate, 0.1% Triton X-100 in molecular-grade water), adjusted to ensure complete sample coverage. Labeled smFISH probes were used at a 1:50 dilution; in cases where signal strength was low, concentrations up to 1:30 were applied. Primary antibodies (as listed above) were included in the hybridization mix. Next, samples were washed twice for a total time of 30 min at 37°C with shaking (200 rpm) in Wash Buffer A. This was followed by incubation with secondary antibodies (Invitrogen, 1:500) and Hoechst 33342 (Invitrogen, 1:3000), both diluted in Wash buffer A, for 1 hour at 37°C, protected from light. Hoechst staining was used to label nuclei. Subsequently, two additional washes were performed with Wash Buffer A for 15 min each. After one final wash in Wash buffer B (2x SSC in molecular-grade water), stained samples were mounted in Vectashield (Vector Laboratories).

### Confocal microscopy

To visualize immunostained MP niches, 12-bit z-stack images were acquired using either a Leica TCS SP8 confocal microscope or a Nikon AX confocal laser scanning microscope (Nikon, Tokyo Japan). The Leica microscope was equipped with PMT detectors and a 40x oil immersion objective (NA 1.3), and the Nikon AX microscope with GaAsP detectors and a 40x silicone oil objective (NA 1.25). Excitation lasers used were 405nm, 488nm, 552nm and 638nm for the Leica, and 405nm, 488nm, 561nm and 640nm for the Nikon AX. Detection windows were set according to the number of channels. Laser power and gain for the Mef2, and Armadillo channels were kept constant within each experiment. To facilitate reproducibility of the plotted raw Mef2 fluorescence intensity values, we provide the specific imaging settings used: averaging of 2, z-step 0.8 µm, dwell time 1.0 (Nikon) /1.2 µsec (Leica). For all other quantifications, intensity values were normalized to an internal control. All images were acquired at 1024×1024 resolution, with a pixel size of 0.285 µm/px (Leica) and 0.31 µm/px (Nikon AX).

To accurately capture smFISH patterns and avoid photobleaching of the sensitive probes, 16-bit images of smFISH samples (*Mef2, zfh1-long, nkd*) were acquired on a Nikon equipped with a Yokogawa confocal spinning disk unit CSU W1 with a 4.2P MP SCMOS camera, using the same 40X silicone oil objective as above. Excitation lasers used were 405nm, 488nm, 561nm, and 647nm. Images were acquired at 2048 × 2048 resolution with a pixel size of 0.162 µm/px and a z-step of 0.8 µm.

### Imaging analysis and data representation

Confocal images were processed using ImageJ/Fiji (https://fiji.sc). Same brightness and contrast values were used for each channel in one experiment. Confocal images were rotated in Fiji such that the anterior side of the MP niche was oriented to the left and the posterior to the right. In addition, for spinning disk images, the tip of the MP niche was aligned to the top of the image. For data representation and quantification, we consistently selected similar z-planes (as highlighted in Fig. 1C) in which all three regions of interest were clearly visible: Mef2low-IFM-MPs (Cut-low), Mef2high IFM-MPs (Cut-low) found anterior from the air sac and the DFM-MPs (Cut- high). Figures were subsequently created in Affinity Designer 2 and Inkscape.

To quantify nuclear Mef2 protein levels in individual MPs within the representative z-plane from immunofluorescence stainings, ten cells from each region were randomly selected in the Hoechst channel, and their Mef2 signal was traced using the “Freehand selections” tool in Fiji. Cells lacking detectable Mef2 signal, such as epithelial cells, were excluded to ensure that ten Mef2-positive nuclei were analyzed per region. The outlines of the nuclei were saved in the ROI (region of interest manager) manager, and the mean pixel intensity for each cell was measured in Fiji. These values were used to calculate and plot the average Mef2 intensity per region. Note, the Mef2 antibody showed partially high background signal in the epithelium, which made background subtraction as a normalization step unfeasible. However, by using consistent imaging settings (especially laser power and gain), we obtained reproducible and comparable staining across samples.

To quantify *zfh1-long* RNA levels, confocal images were analyzed by first selecting the representative z-plane in the Mef2 channel, which was then used for *zfh1* RNA quantification. In Fiji, the image was converted into 8-bit, and a manual threshold was applied to generate a binary mask that included only the bright dots in the *zfh1- long* RNA channel. The threshold was adjusted based on the overall brightness of each stained sample but was kept within a similar range between control and knockdown conditions. The “Analyze particles” command was then used to detected individual dots (size ranging: 0.1 – 6 μm^2^) in the ROI manager. The number of ROIs were then normalized to the MP area, which was traced and measured in the same z-plane in the Mef2 channel using the “Freehand selections” tool.

To quantify the *Armadillo* knockdown efficiency, Arm protein levels were measure in immunostained samples. The representative single z-plane confocal image was selected from both control and *armadillo* knockdown sample. The MP area was outlined using Fiji’s „Freehand Selections” tool based on Mef2 staining, and the mean Armadillo intensity was measured.

### Micro-CT thorax preparation and scanning

To evaluate the overall effects of *Mef2* overexpression in the Mef2-low IFM- MPs on adult flight muscles, adult thoraces from one-day-old female flies were dissected in 1x PBS by removing the head, wings, legs and abdomen. The samples were submerged in a tube containing 1x PBS for up to 30 min to allow them to settle at the bottom. Occasionally, a few drops of 0.3% PBST were added to reduce the surface tension. The thoraces were fixed with 4% PFA for 3 hours (in the dark, rotating) and washed twice with 1x PBS for 15 min each. Prior to micro-CT scanning, the thoraces were washed with water for 15 min and stained with buffered Lugol’s solution (3.75% Lugol and 0.1 M Sorensen’s buffer) over night. Flies were mounted in 2% low-melting agarose in PCR tubes and scanned on a SkyScan 1272 system (Bruker, Massachusetts, USA) by the Electron Microscopy Core Facility of Heidelberg University. For irradiation, an L10101 microfocus X-ray source (Hamamatsu Photonics) was used with power settings between 50–80 kV and 50–70 µA. Scans (16-bit) were acquired with a CMOS 16 MP camera (Photonic Science) with an image pixel size of 1.5 µm, 0.2° rotation steps, a frame average of 6–10 and random movement correction set between 5–15. The scanning was performed with 180° rotation. Reconstruction of the acquired scans was done with the NRecon software (Bruker, v2.2.0.6). Reconstruction parameters such as misalignment compensation, beam hardening, and ring artifact correction were set individually for each scan to achieve optimal results. Images were processed using Dragonfly software (version 2022.2; available at https://www.theobjects.com/dragonfly) on available workstations at the Nikon Imaging Center of Heidelberg University. Three representative slice views were selected by navigating through the 3D microCT volume. Brightness and contrast settings were adjusted as needed to enhance tissue contrast and highlight anatomical features.

### scRNA-seq data integration and analysis

To investigate *Mef2* heterogeneity at the transcriptional level in the MP niche, scRNA-seq datasets from previously published studies were used (Bageritz et al., 2019; Everetts et al., 2021). 10x Genomics data from three replicates (one from Bageritz et al., two from Everetts et al.) were analyzed using the Cell Ranger pipeline (version 7.1.0). Sequencing reads were aligned to the *Drosophila melanogaster* reference genome (BDGP6 v105 (GCA 000001215.4)). Downstream analysis, including integration of the datasets, was performed using the R package Seurat (version 5.0.1), following the Satija Lab’s guidelines (https://satijalab.org/seurat/). High-quality cells were retained by filtering for those with a minimum of 200 detected genes and a total transcript count greater than 2,000. The percentage of mitochondrial transcripts in all replicates was below 10%, so a cutoff was not applied. The anchor-based CCA integration method was used to align the three replicates. MP cells were extracted from the integrated dataset based on the expression of known MP markers (Bageritz et al., 2019). A second round of clustering on the MP subset revealed a group of cells co-expressing MP and epithelial markers, suggesting the presence of cell doublets. These cells were filtered out, resulting in a final set of around 13,000 high-quality MP cells. Initial clustering was strongly influenced by cell cycle differences within the MPs. To address this, cell cycle scores were calculated separately to each replicate using a manually curated list of cell cycle-related genes previously identified as differentially expressed in the Everetts dataset (Everetts et al., 2021). Cell cycle regression was then performed as outlined in the *Cell-Cycle Scoring and Regression* vignette of the Satija laboratory. To identify differentially expressed genes between the Mef2-low and Mef2-high expressing IFM-MPs, Seurat’s FindConservedMarkers function was used. The resulting gene list was filtered to retain only genes with an average log2 fold change of ≥ 0.7 of across all three replicates.

### Data representation and statistical test software

All data analysis, statistical testing, and plot generation were performed in R. As the data did not follow a normal distribution, as assessed by Shapiro-Wilk-test, all group comparisons were performed using the Wilcoxon rank-sum test. Data visualization was carried out using ggplot2 package. Statistical significance was indicated as follows: P < 0.05 = *, P < 0.01 = **, and P < 0.001 = ***.

## Acknowledgments

We are grateful to Hadi Boukhatmi, Aurelio Teleman, and Julia von Maltzahn for providing comments on the manuscript. We thank Ulrike Engel and Christian Ackermann from the Nikon Imaging Center, and Réza Shahidi and Larissa Eis from the Electron Microscopy Core Facility of Heidelberg University for providing support. We are grateful to Magdalena Schlotter for helping to set up the smFISH protocol in the lab. We are grateful to the fly community for providing fly stocks and reagents, in particular Hadi Boukhatmi for the Enh3-Gal4, UAS-GFP fly line, Ruth Lehmann for providing the Zfh1 antibody, and Katrin Domsch for sharing the Twist and Mef2 antibody. This work was supported by the Deutsche Forschungsgemeinschaft (DFG, German Research Foundation) – project number 331351713 – SFB1324 on “Mechanisms and Functions of Wnt Signaling”, and by the Health + Life Science Alliance Heidelberg Mannheim and received state funds approved by the State Parliament of Baden-Württemberg. The authors acknowledge support by the state of Baden-Württemberg through bwHPC.

