## Supplementary information for "Gradient of Wnt signaling facilitates Mef2 heterogeneity and limits commitment of the developmental muscle progenitor pool"

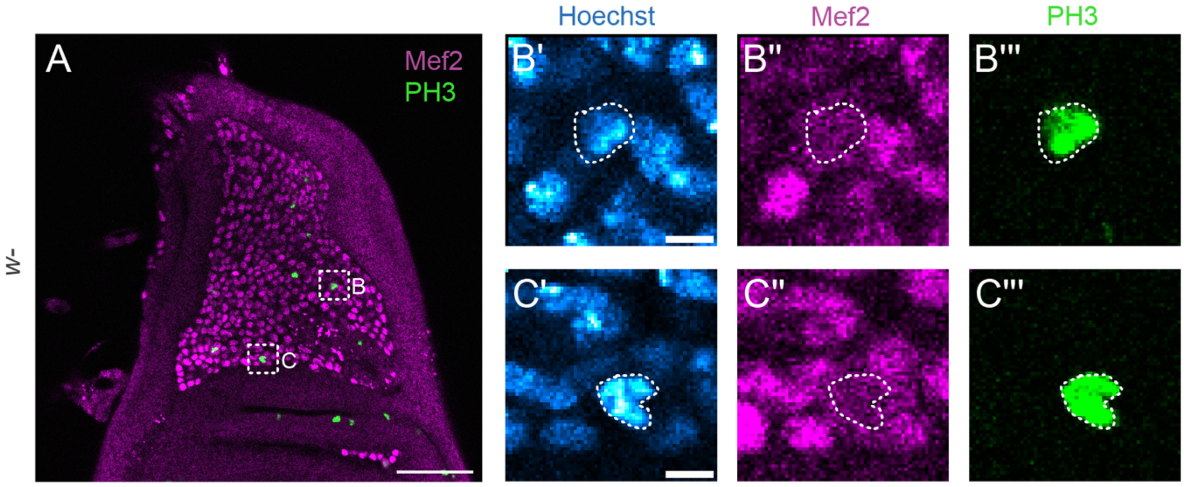


**Supplementary Figure 1: Mitotic muscle progenitors in Mef2-high regions exhibit reduced Mef2 level. A)** Single z-plane confocal image of the larval muscle progenitor (MP) niche stained for Mef2 and Phospho-histone H3 (PH3) to label mitotic cells. Scale bar: 50 µm. Close-ups of the Mef2-high IFM-MP **(B-B''')** region and the DFM-MP **(C-C''')** region showing PH3-positive MPs with reduced Mef2 signal. Scale bar for close-ups: 4 µm.

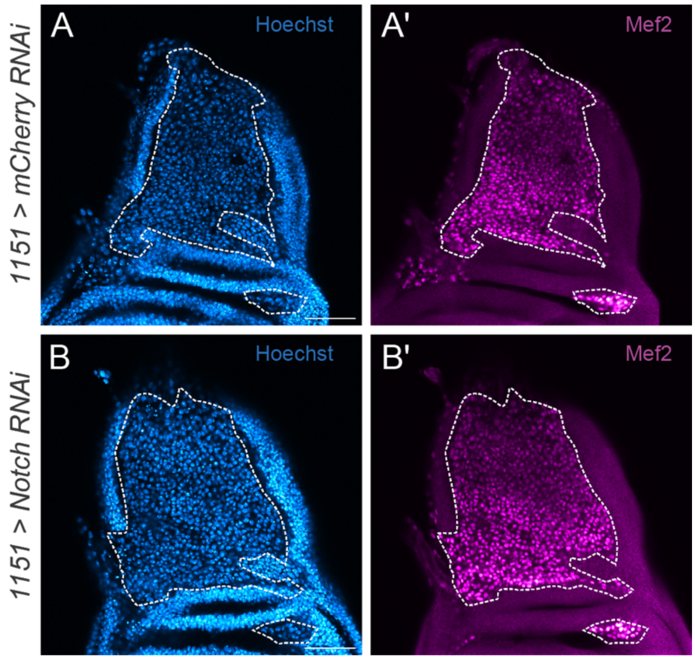


**Supplementary Figure 2: *Notch* perturbation in the muscle progenitor pool does not affect Mef2 heterogeneity. A–B)** Representative single-plane confocal images of the larval muscle progenitor niche in control **(A, A')** and *Notch* RNAi **(B, B')** conditions using the MP-specific 1151-Gal4 driver (n=6). Hoechst staining marks all nuclei **(A, B)**. Mef2 protein, detected by immunofluorescence **(A', B')**, shows the heterogeneous expression pattern in control. In the *Notch* knockdown condition, the spatial heterogeneity of Mef2 is maintained. The MP region is outlined by a dashed line. Scale bar: 50 µm.

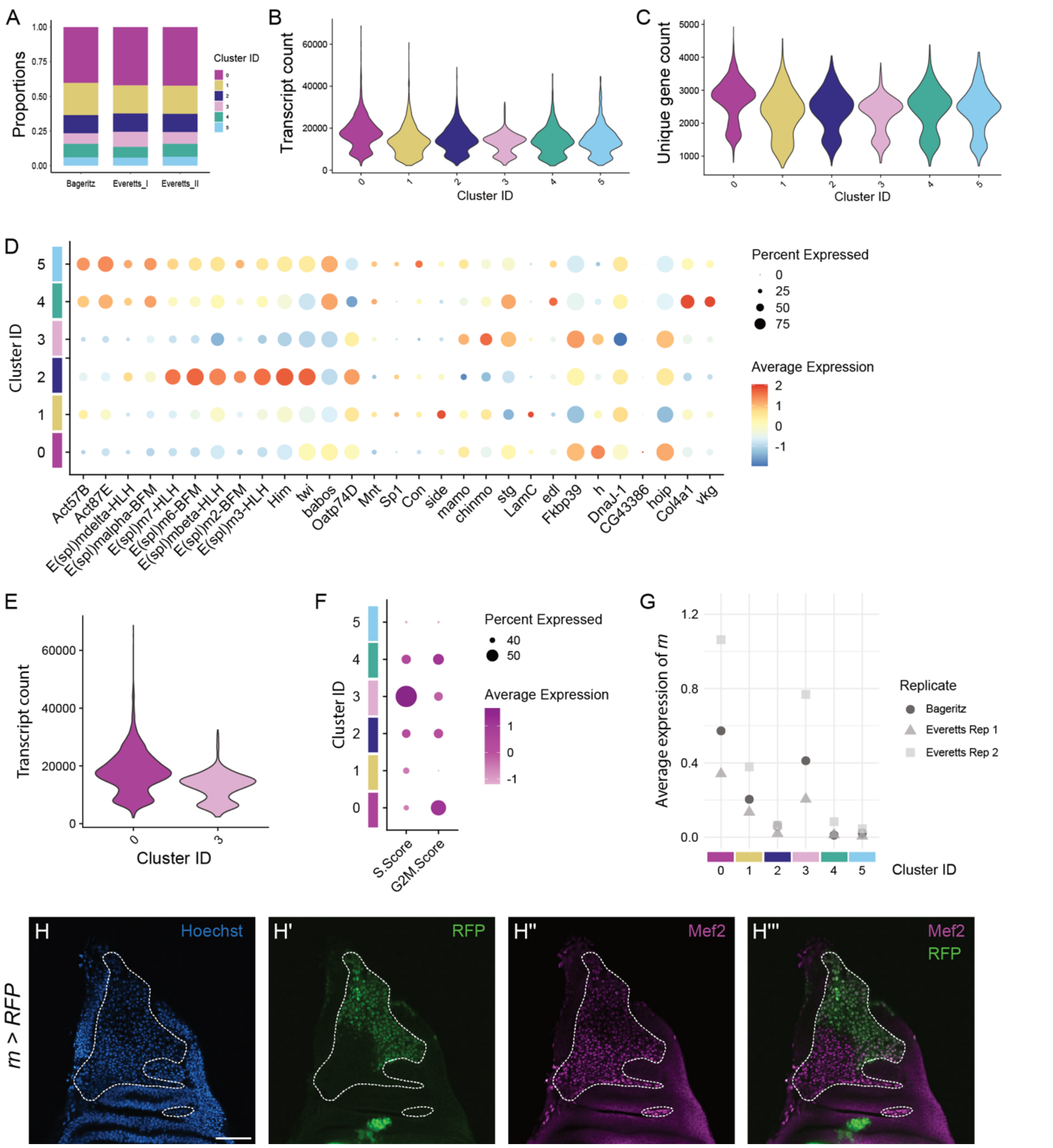

**Supplementary Figure 3: Integration of single-cell RNA-sequencing datasets validates and characterizes Mef2-low muscle progenitor population. A)** Contribution of the three biological replicates to each cluster in the integrated dataset. All replicates contribute comparably to the identified clusters, supporting successful dataset integration. **B)** Total transcript counts and **C)** unique gene count per cluster. **D)** Dotplot cross-referencing our integrated dataset with marker genes from Zappia and colleagues (Zappia et al., 2020). **E)** Total transcript count comparison between the Mef2 low-clusters 0 and 3. **F)** Cell cycle phase scoring reveals residual differences between clusters 0 and 3 in S and G2/M scores, despite regression of cell cycle effects. **G)** Average expression of the Mef2-low marker gene *rotund* (*rn*) across all clusters and replicates. **H)** Representative single-plane confocal image showing rn-Gal4 > UAS-RFP expression in the dorsal region of the MP niche, overlapping with the Mef2-low area identified by immunostaining (n=4). The MP region is outlined by a dashed line. Scale bar: 50 µm.

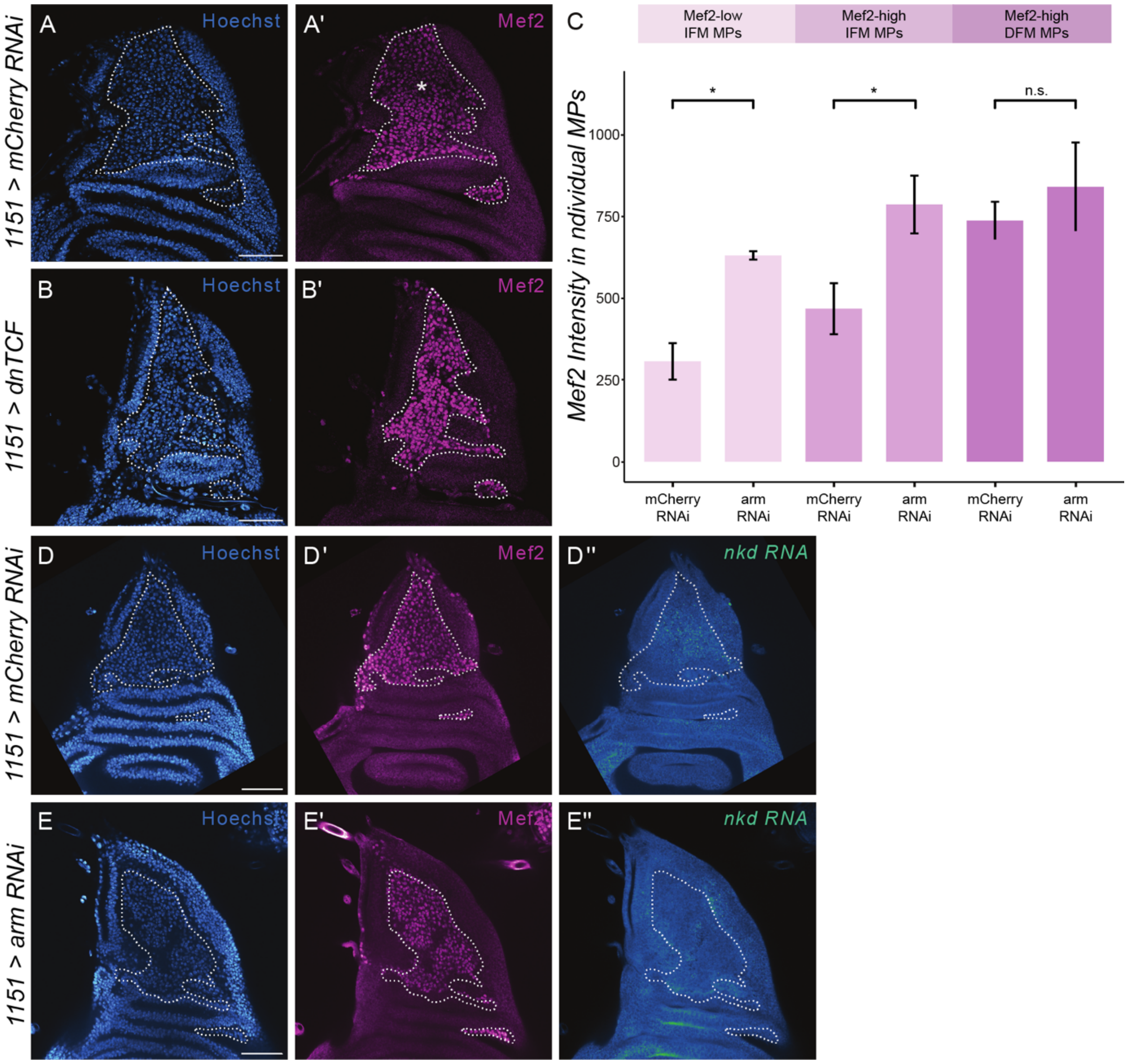


**Supplementary Figure 4: Mef2 regulation is mediated through the Arm–TCF axis of Wnt/β-catenin signaling. A–B)** Representative single-plane confocal images of the larval muscle progenitor (MP) niche in control **(A)** and dominant-negative TCF (dnTCF; **B**) conditions, using the MP-specific 1151-Gal4 driver. Dashed lines outline the MP region. Expression of dnTCF phenocopies *arm* knockdown, resulting in elevated Mef2 levels. (Control: n=5; dnTCF: n=3). **C)** Quantification of Mef2 protein levels in three anatomically defined areas (Mef2-low IFM-MP, Mef2-high IFM-MP and Mef2-high DFM-MP) following misexpression of dnTCF. Mef2 intensity is significantly increased in the Mef2-low and Mef2-high IFM-MP regions. Wilcoxon statistical test was performed. (Control: n=5; dnTCF: n=3). **D–E)** Representative single-plane confocal images of the larval muscle progenitor niche following *arm* knockdown. Hoechst staining marks all nuclei (**D, E**); Mef2 protein is detected by immunofluorescence **(D', E')**; and *nkd* mRNA is visualized by smFISH (D'', E''). A reduction in *nkd* transcript level is observed (Control: n=7; *arm* knockdown: n=7, across two independent experiments). Scale bar: 50 µm.

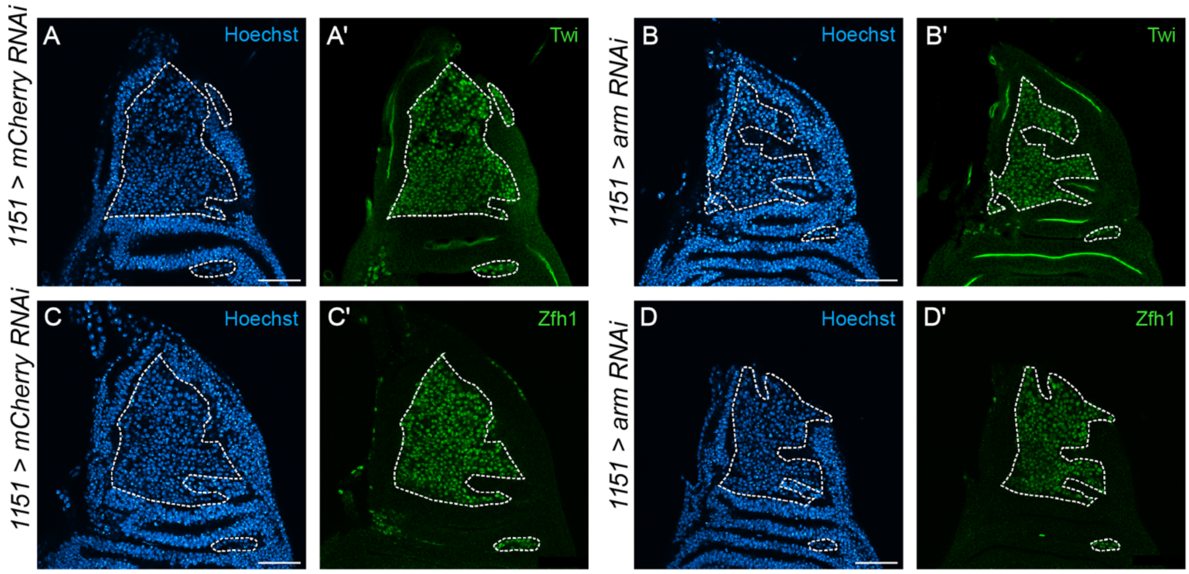


**Supplementary Figure 5: Wnt/β-catenin signaling perturbation does not alter Twist or Zfh1 levels in the Mef2-low region at the L3 larval stage. A-B)** Representative single-plane confocal images of the larval muscle progenitor niche stained for Twist (Twi) in control (**A**, n=6) and *arm* knockdown (**B**, n=6) conditions using the MP-specific 1151-Gal4 driver. Twi protein levels and spatial distribution show no detectable changes in the Mef2-low domain following *arm* knockdown. Scale bar: 50 µm. **C-D)** Representative single-plane confocal images of Zfh1 staining in control (**C**, n=7) and 1151-Gal4-driven *arm* knockdown (**D**, n=7) in the larval muscle progenitor niche. Zfh1 protein abundance or spatial distribution appear unchanged in the Mef2-low IFM-MP domain following *arm* knockdown. The MP region is outlined by a dashed line. Scale bar: 50 µm.

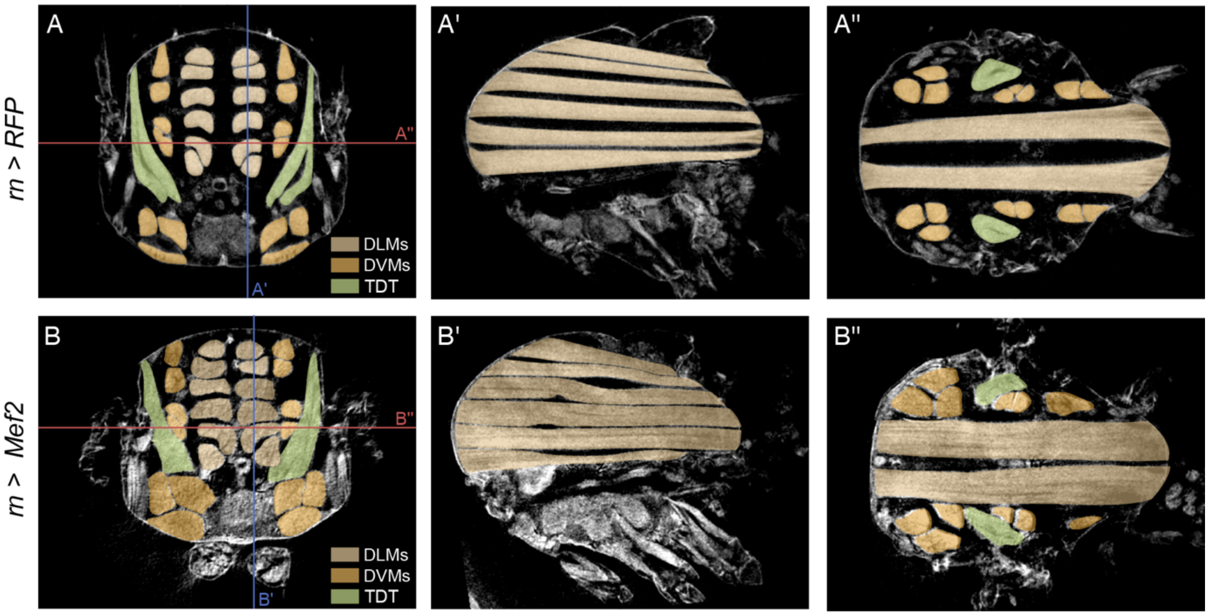


**Supplementary Figure 6: Micro-CT analysis reveals no change in DLM or DVM fiber number upon *Mef2* overexpression in Mef2-low muscle progenitors. A–B)** Representative Micro-CT images of adult female thoraces in control (rn-Gal4 > UAS-RFP; **A**, n=3) and *Mef2* overexpression (rn-Gal4 > UAS-Mef2; **B**, n=3) conditions. Red and blue lines indicate the levels of the two cross-sectional views shown **A'–A'', B'–B'',** respectively) Cross-sectional views at two different planes illustrating the organization and fiber number of the dorsal longitudinal muscles (DLMs, light orange) and dorsal ventral muscles (DVMs, dark orange). The tergal depressor of the trochanter (TDT, or jump muscle) is marked in green. No difference in DLM or DVM fiber number is observed between control and *Mef2* overexpression conditions.
